# Delayed effect of drought on the xylem vulnerability to cavitation in *Fagus sylvatica* L

**DOI:** 10.1101/2020.05.14.096347

**Authors:** Stephane Herbette, Olivia Charrier, Hervé cochard, Têté Severien Barigah

## Abstract

Knowledge on variations of drought resistance traits is needed to predict the potential of trees to adapt to severe drought events expected to be more intense and frequent. Xylem vulnerability to cavitation is among the most important traits related to drought-induced mortality and exhibits a large variability between species. Acclimation of this trait to environmental conditions implies changes in the xylem structure and organization, leading previous studies to investigate its variations under conditions preserving growth. In European beech saplings, we assessed the effect of droughts of on the vulnerability to cavitation in branches that develop during recovery. The newly formed branches displayed lower vulnerability to cavitation in the plants that underwent the severest droughts leading to native embolism; the pressure that induces 50% loss of conductance being of −3.98 MPa in severely droughted plants whereas it was of −3.1 MPa in control plants, respectively. Although unexpected, these results argue for an acclimation, and not a weakening, of this trait to drought events.

**Key message:** severe water stress make the future developed shoots less vulnerable to xylem cavitation.

## Introduction

In a context of climate changes, the study of variations among and within species of drought resistance mechanisms is needed to predict their potential of adaptation to drought events expected to be more intense and frequent. For woody species, drought-induced dieback is more likely due to xylem hydraulic failure (Anderegg et al. 2015; 2016) caused by the cavitation events in the xylem conduits, even if the carbon starvation can also contribute to this dieback (Hartmann 2015). A recent model (Martin-St Paul et al. 2017) has pointed out the importance of the coordination between the water potential causing stomatal closure and the xylem pressure that provokes cavitation, i.e. xylem vulnerability to cavitation (VC) (Martin-St Paul et al. 2017). This VC is considered as a critical trait for drought tolerance in woody (Choat et al. 2012; Lens et al. 2013) and in herbaceous species (Lens et al. 2016; Volaire et al. 2018). Moreover, a global analysis pointed out the narrow hydraulic safety margin at which woody species commonly operate (Choat et al. 2012), inferring that research is needed on the variability for xylem VC under the current climate changes. Whereas a high variability of VC was reported between species in several studies, ranging from more than −1 to −19 MPa (Choat et al. 2012; Larter et al. 2015; Martin-St Paul et al. 2017), the extent of variation within species has been reported to be considerably less, in a range usually lower than 1 MPa (e.g. Herbette et al. 2010). However, the studies are rather few, and knowledge is lacking on the drivers of within species variability for VC, and thus on the extent of this variability.

The genetic variability for VC is rather limited, in both natural populations (Wortemann et al. 2011; Lamy et al. 2011) and cultivated genotypes (Jinagool et al. 2015; 2018). An uniform selection has been reported to explain the weak variability for VC in a pine species (Lamy et al., 2011). The within species variation in VC is largely attributed to environmental factors (Martinez-Vilalta et al. 2009; Herbette et al. 2010; Worteman et al. 2011). These factors include sunlight exposure (Barigah et al. 2006; Herbette et al. 2010), nutrient availability (Harvey and Van Den Driessche 1999; Plavcová and Hacke 2012), and mostly soil water contents (Awad et al. 2010). Acclimation of xylem VC to environmental conditions implies changes in the xylem structure and organization. This trait relies on conduits properties from intervessel pit to xylem structure (Lens et al. 2013). Changes in xylem anatomy can occur only in the growing period during wood formation, as xylem conduits are made of dead cells with no short-term capacity to acclimatize or adjust to changes in hydraulic demand. For example, no change in VC was observed for beech branches across contrasted seasons along a growing season, or for branch samples put in humid and dark conditions during several weeks (Herbette et al. 2010). A decrease in stem VC related to drier conditions has been observed during the growing period, providing that the growth is maintained (Awad et al. 2010; Fichot et al. 2010; Plavcová and Hacke 2012). Yet, the climate change models predict further increases in the frequency and the severity of extreme drought events (IPCC 2013), and such drought events occur mainly during the dry season when there is few or no more growth, and if there is, it stops the growth. Knowing the effects of severe drought conditions on the VC, as well as on other drought-resistance mechanisms, may help us to predict the response of woody species impact to climate changes.

In this study, we assessed the impact of different duration of water deprivation on VC of branches developed after resumption from water shortage. The ‘‘Cavitron” centrifugal technique (Cochard et al. 2005) is an effective, rapid method for large-scale investigation of VC (Lamy et al. 2011; Wortemann et al. 2011; Cochard et al. 2016), especially for beech samples (Herbette et al. 2010; Wortemann et al. 2011; Barigah et al., 2013 Stojnic et al. 2017), and could thus be used in this study. We carried out the experiment on European beech *(Fagus sylvatica* L.) saplings with water stress applied at the end of spring i.e. after the growth has stopped. It is a widespread species of the European forest and known to be sensitive to drought (Bréda et al.2006). We thus assumed that a severe drought would make the new branches more vulnerable to cavitation as the carbon reserve would be impaired and thus would impact the development of new axes. Indeed, it is well established that there is a carbon cost involved in developing cavitation-resistant xylem (Hacke and Sperry 2001).

## Materials and Methods

### Plant material and growth conditions

We carried out the experiment in a greenhouse at the INRA research station of Clermont-Ferrand in France (45877’N. 3814’E; altitude of 300 m) during 2008 and 2009. Plant material and growth conditions were previously described in Barigah et al. (2013). We excavated from a nursery stand and transplanted saplings of European beech *(Fagus sylvatica* L.) of two-years old in November 2007 into 20 liters pots filled with a 1:2 (v/v) mixture of peat and local forest soil. The 159 saplings grew up in a greenhouse with daily irrigation to field capacity (FC) from planting to the beginning of the water-stress treatment. We set up a drip irrigation controlled by a timer to supply the plants with tap water. We applied a severe water-stress to 144 individuals from June 30^th^ 2008 by ceasing watering them for 1 to 12 weeks (12 individuals per week treatment), except to a set of 15 control plants we watered at field capacity. At the end of each of the 12 drought periods, the plants were re-watered to FC after data collection completed (Barigah et al. 2013). Then, we kept all the plants at FC after the resumption from the water shortage, in the same growth conditions than before the drought. While most of the plants were used for sampling in Barigah et al. (2013) or died because of the drought severity, 6 plants per treatment were kept from any cutting and recovered in 7 treatments: control and drought duration from 1 to 6 weeks. Among these latter, 4 to 6 plants per treatment developed new branches of at least 0.3m long after the drought that were harvested on July 2009 for measuring their VC.

### Stomatal conductance and native embolism

Leaf conductance to water vapour (*g*_s_) was measured each week during the drought treatment with a steadystate porometer (Li-1600, Li-Cor, Lincoln, Ne, USA) equipped with a broad leaf chamber. Measurements were performed on leaves from 5 to 40 different individual saplings per water treatment. Minimal *g_s_* was measured between 12:00 and 14:00 solar time on sunny days. Depending on the climatic conditions when measuring g_s_, we could observe variations in data, leading us to increase the number until 40 individual saplings per condition.

Native embolism measurements were detailed in *Barigah* et al. (2013).

### Vulnerability to cavitation

Intact stems of at least 0.3m long were defoliated and then harvested from 4 to 6 individual saplings for each drought duration. Stems were wrapped in moist paper, placed in plastic bags to avoid dehydration, and cold-stored (4 °C) for not more than 7 d before analysis of their VC. Segments of 0.28 m-long were recut using a razor blade to refresh the ends, just before the measurement of their VC by using the Cavitron technique (Cochard et al. 2005). Xylem pressure (*Ψ*_x_) was first set to a reference pressure (−0.5 MPa) and the sample maximal conductance (*K*_max_) was determined. The xylem pressure was then set to a more negative pressure for 2 min and the new conductance (*K*) was determined. The procedure was repeated for more negative pressures (with −0.25 or −0.50 MPa increments) until PLC reached at least 90%. Vulnerability curves were generated for each species as plots of PLC versus Ψ_x_. Data were then fitted with the following exponential–sigmoidal functions:

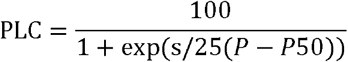

Where *P*_50_ is the pressure causing 50% loss of conductance and *s* is the slope value at this point.

### Model simulations

To simulate the impact of the plasticity of P50 on plant drought resistance, we used the mechanistic SurEau model (Martins St-Paul et al 2017, Cochard 2019) to predict the timing of hydraulic failure under standardized conditions. The model was parametrized to match as much as possible the characteristics of the plant material used in this study and the environmental growth conditions. In more details, the key plant parameters of the simulation were the following: leaf area: 0.1 m^2^; shoot length: 0.6 m; basal diameter: 0.6 cm; maximum stomatal conductance: 300 mmol/s/m^2^; minimum cuticular conductance: 2 mmol/s/m^2^; Leaf Π_0_: −2.0 MPa; maximum whole plant conductance: 0.28 mmol/s/m^2^/MPa. The P50 and s parameters were identical for all the plant organs, and P50 was allowed to vary between −3 to −4MPa (see results). The environmental conditions were set as follow: night/day air temperature: 15/25°C; night/day RH: 80/40%; PAR: 650 μmol/s/m^2^; wind speed: 1m/s; soil volume: 20 dm^3^; available soil water content: 7.2 kg. We assumed that stomata closed during water stress in response to changes in leaf turgor pressure (Martin St-Paul et al 2017). The stimulation started at midnight with a soil at full water capacity and the plant were let to transpired until 90% of embolism was formed in the stem, that was considered as the point of hydraulic failure.

### Statistics

An analysis of variance (ANOVA) was used to test the effects of watering regimes on VC. Means were compared with Tukey’s multiple range with P-value < 0.05, after checking homogeneity and normality of the data. All the measured and derived data were subjected to the statistical analysis using a software package (XLSTAT v7.5.3, Addinsoft, Paris, France).

## Results and discussion

The beech sapling were submitted to a drought by ceasing watering for up to 12 weeks. The stomatal conductance declined starting from the first week of the drought, then it reached very low and constant values after 3 weeks (fig. 1), indicating a stomatal closure starting at this time. We could not measure the *g*_s_ beyond 7 weeks of drought as the leaves dessicated. The rate of native embolism started to increase 3 weeks after the water-stress inception and reached their maximal values starting from the 9^th^ week (fig. 1). After each week of drought, a set of plants were rewatered to full capacity up to the following year. This experimental device allows assessing the delayed effects of a wide range of water stress from the weak decrease in *g*_s_ to the death, through the stomatal closure point and the accumulation of embolism.

**Fig. 1.**
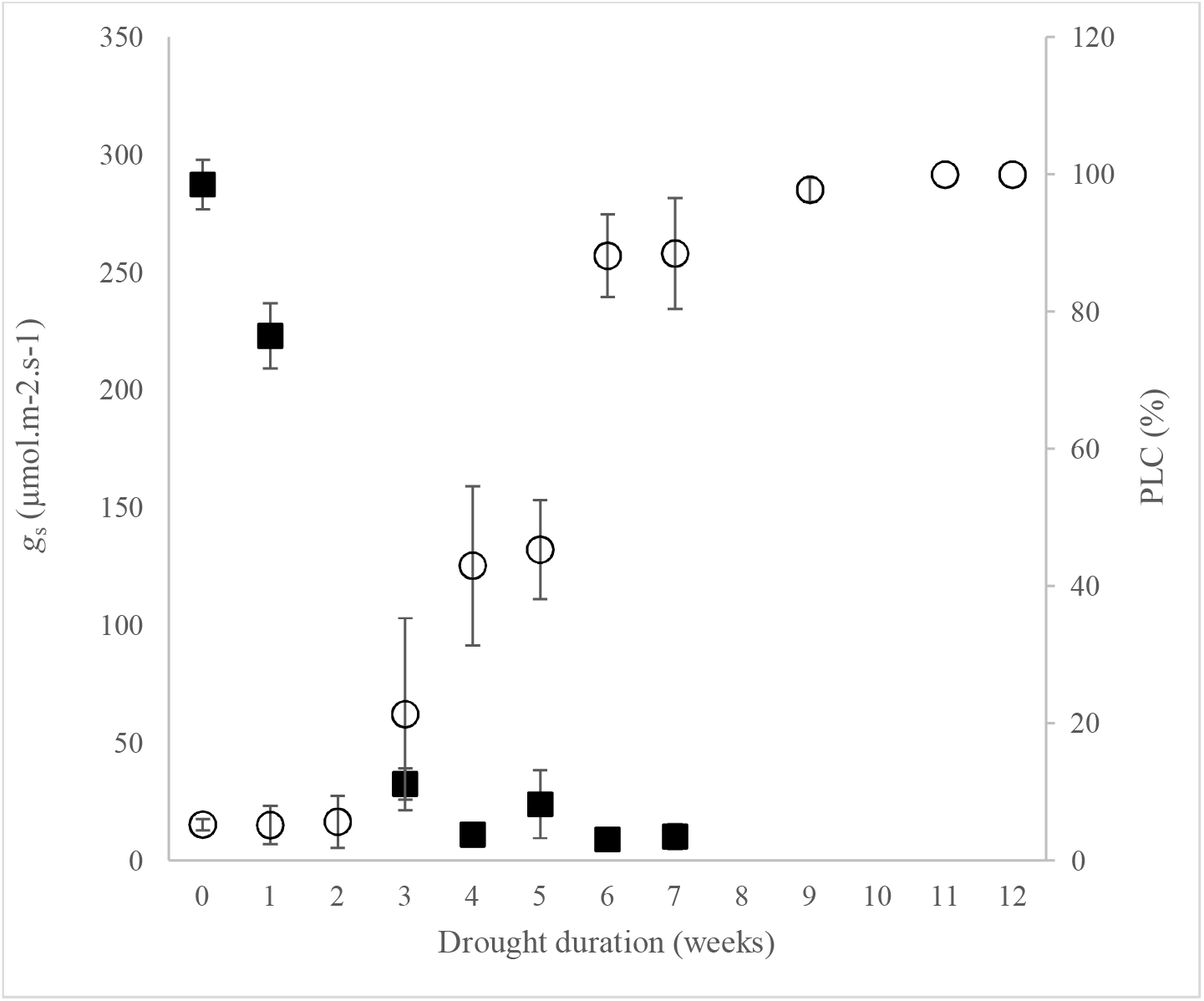
Evolutions of leaf stomatal conductance *g*_s_) and embolism level (PLC) during the drought. Surveys of *g*_s_ (black square) and PLC (open circle) were carried on leaves and stems, respectively, of beech saplings that underwent a drought by irrigation stop up to 12 weeks. No *g*_s_ could be measured beyond 7 weeks as the leaves were completely dry. PLC data were from Barigah et al. (2013). Data are mean values (n = 5 to 40 for *g*_s_ and n = 6 or 7 for PLC) and errors bars are S.E.

We selected plants that recovered and that were not previously cut for sampling. All plants were let to finish their growth period, with watering at field capacity, to ensure that there is no delay in growth between sets of plants. Then, VC was measured on branches that newly formed under watering at field capacity. For control plants, the VC increased from −3.43 MPa (+/- 0.17) just before the drought experiment to −3.10 MPa (+/- 0.30) in newly formed stems in the following year. Such difference can be explained by difference in growth conditions and physiological status of the plants between the 2 years. To avoid such effects, the drought effect was analyzed by comparing newly formed stems of the different treatments.

The figure 2 shows the P50 mean values for these branches depending on the drought duration the plants underwent on the previous year. The P50 decreased from −3.10 to −3.98 MPa for 6weeks of drought, and mean values became significantly different from control plants after 4 weeks of drought. Our results clearly demonstrate that beech sapling developed new branches with lower VC after the plants underwent severe drought condition that leads to hydraulic failure. The lowest VC was obtained after 6 weeks of drought and no death was observed in this condition (Barigah et al. 2013). Thus, a selection effect of the drought can be rule out and led us conclude that the change in VC was only due to phenotypic plasticity. We could not analyzed the VC for longer drought periods as part or most of the plants died (Barigah et al. 2013). Our results would explain how marginal beech populations have a lower VC and are drought-tolerant (Stojnic et al. 2017; Bolte et al. 2017), whereas there is little or no difference in VC between core populations. Indeed, marginal populations in drier conditions are more prone to undergo severe water deficit that would induce a decrease in VC. The extent of this decrease in *P*_50_ is rather large (0.88 MPa or 28% of the control value) when compared to variations observed for this species due to other environmental conditions (Barigah et al. 2006; Herbette et al. 2010), and would have a significant impact on the drought resistance of this species. We tested this impact using the hydraulic soil-plant model proposed by Martin-St Paul et al. (2017) and Cochard (2019) that predicts the time to plant hydraulic failure. The model parameters were first turned to match the dynamics of PLC and *g*_s_ shown in figure 1 and considering a *P*_50_ of −3.1 MPa. We then ran simulations with the same set of parameters, allowing only *P*_50_ to change according to the values reported in figure 2. Simulations showed that a decrease of *P*_50_ from −3.10 to −4 MPa would delay by 17 days the timing of hydraulic failure, and thus shoot mortality (fig. 3). The next step will be to assess the delayed effect of drought on several drought-related mechanisms, in order to investigate the potential of adaptation to drought events using the same model.

**Fig. 2.**
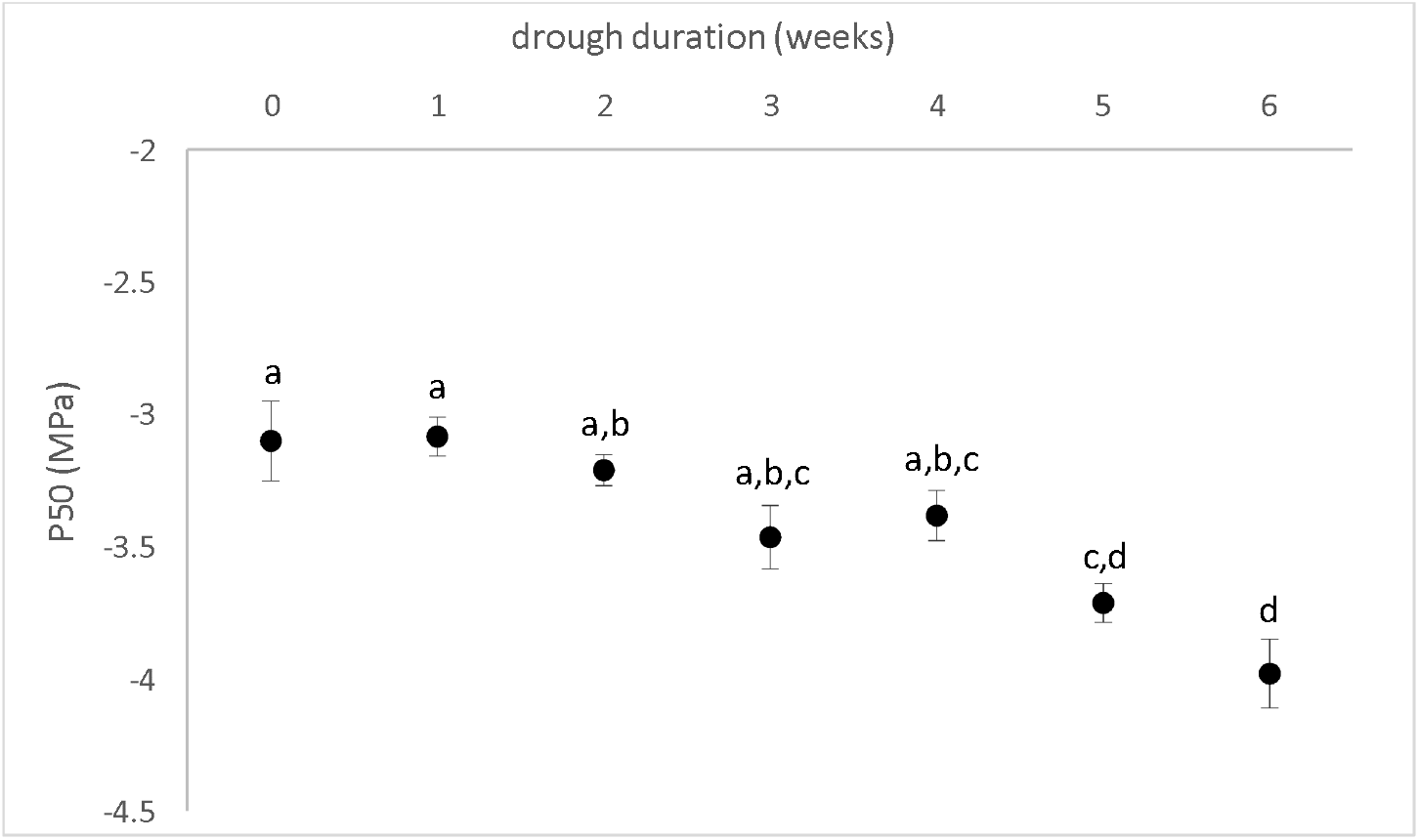
Vulnerability to cavitation (*P*_50_) of branch developed after droughts of different duration. Vulnerability to cavitation was measured on branches that developed the year after the beech saplings underwent a drought by withholding up to 6 weeks. No data were recorded for saplings assigned to more than 6 weeks of water stress since some or most of the plants did not recover. Data are mean values (n = 4 to 6) and errors bars are S.E. Data were analyzed using ANOVA tests and Tukey’s tests (P < 0.05). Different letters indicate significant differences for P50 between the drought spans.

**Fig. 3.**
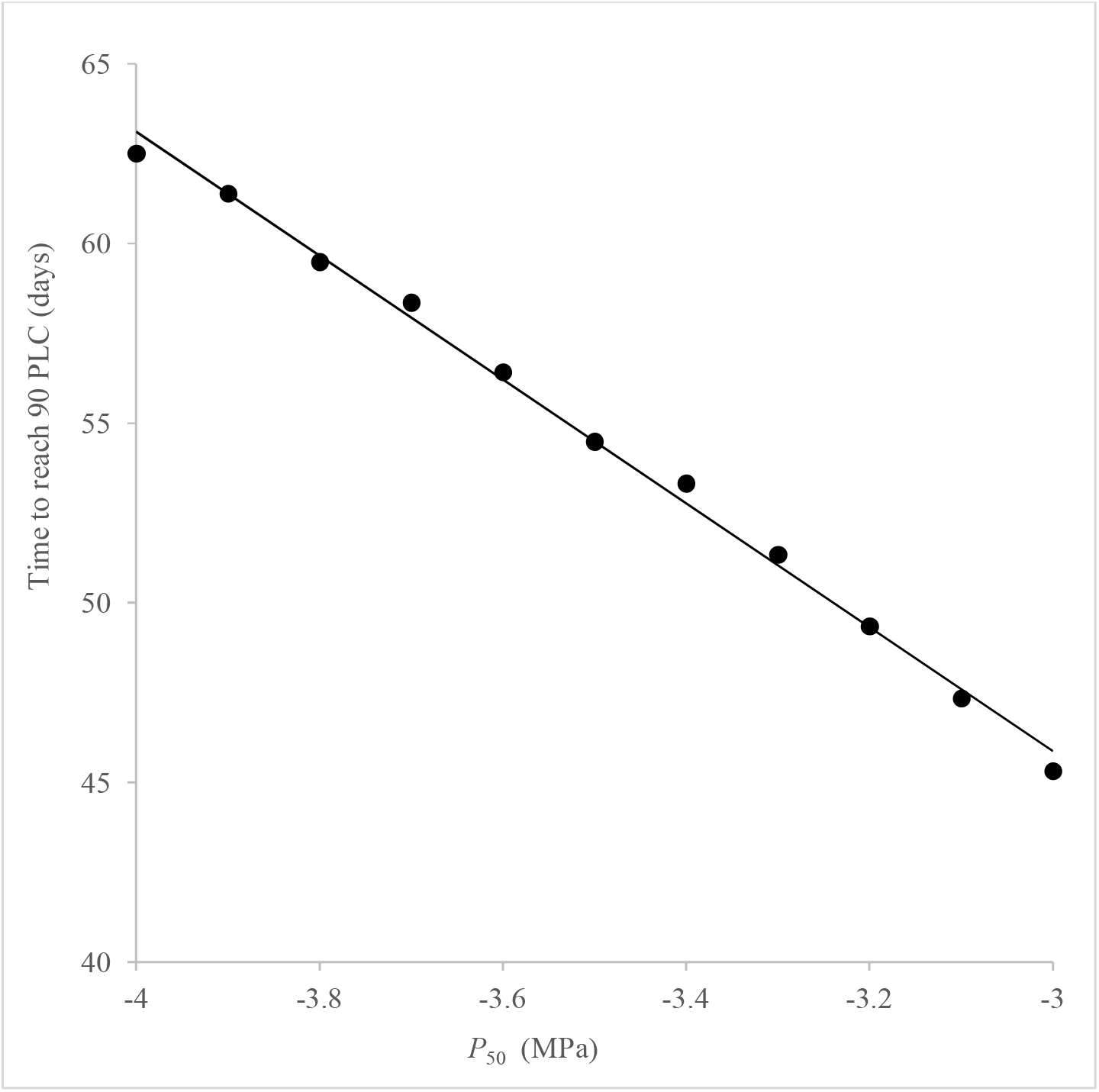
Model simulation of the time to reach hydraulic failure for the range of VC observed in this study. The model developed by Martin St Paul et al. (2017) was used to estimate the time to reach 90 % loss of conductance (leading to death according to Barigah et al. 2013) depending on the VC the branch developed in this study. With the exception of the changes in *P*_50_ made to satisfy the hypothesis tested, all other parameters were kept constant.

This drought-induced decrease in VC is rather surprising, as we assumed that drought would increase the VC as the carbon assimilation is impaired. The droughted plants had a lower growth (data not shown) when compared to control, suggesting that the acclimation for VC was prioritized for carbon allocation over the growth. The challenge will be to elucidate the mechanism underlying this delayed effect, especially because this acclimation occurs in newly formed branches that did not underwent the drought.

## Conflict of interest

The authors declare that they have no conflict of interest.

## Notes

### Competing Interest Statement

The authors have declared no competing interest.

## References

Anderegg WR, Klein T, Bartlett M, Sack L, Pellegrini AF, Choat B, Jansen S (2016) Meta-analysis reveals that hydraulic traits explain cross-species patterns of drought-induced tree mortality across the globe. Proc Natl Acad Sci U S A 113: 5024–5029

Anderegg WRL, Flint A, Huang C-Y, Flint L, Berry JA, Davis FW, Sperry JS, Field CB (2015) Tree mortality predicted from drought-induced vascular damage. Nature geoscience 8: 367

Awad H, Barigah T, Badel E, Cochard H, Herbette S (2010) Poplar vulnerability to xylem cavitation acclimates to drier soil conditions. Physiol Plant 139: 280–288

Barigah TS, Charrier O, Douris M, Bonhomme M, Herbette S, Ameglio T, Fichot R, Brignolas F, Cochard H (2013) Water stress-induced xylem hydraulic failure is a causal factor of tree mortality in beech and poplar. Ann Bot 112: 1431–1437

Barigah TS, Ibrahim T, Bogard A, Faivre-Vuillin B, Lagneau LA, Montpied P, Dreyer E (2006) Irradiance-induced plasticity in the hydraulic properties of saplings of different temperate broad-leaved forest tree species. Tree Physiol 26: 1505–1516

Bolte A, Czajkowski T, Cocozza C, Tognetti R, de Miguel M, Psidova E, Ditmarova L, Dinca L, Delzon S, Cochard H, Raebild A, de Luis M, Cvjetkovic B, Heiri C, Muller J (2016) Desiccation and Mortality Dynamics in Seedlings of Different European Beech *(Fagus sylvatica* L.) Populations under Extreme Drought Conditions. Front Plant Sci 7: 751

Bréda N, Huc R, Granier A, Dreyer E (2006) Temperate forest trees and stands under severe drought: a review of ecophysiological responses, adaptation processes and long-term consequences. Ann For Sci 63: 625–644

Choat B, Jansen S, Brodribb TJ, Cochard H, Delzon S, Bhaskar R, Bucci SJ, Feild TS, Gleason SM, Hacke UG, Jacobsen AL, Lens F, Maherali H, Martinez-Vilalta J, Mayr S, Mencuccini M, Mitchell PJ, Nardini A, Pittermann J, Pratt RB, Sperry JS, Westoby M, Wright IJ, Zanne AE (2012) Global convergence in the vulnerability of forests to drought. Nature 491: 752–755

Cochard H (2019) A new mechanism for tree mortality due to drought and heatwaves. bioRxiv: 531632.

Cochard H, Damour G, Bodet C, Tharwat I, Poirier M, Ameglio T (2005) Evaluation of a new centrifuge technique for rapid generation of xylem vulnerability curves. Physiol Plant 124: 410–418

Cochard H, Torres-Ruiz JM, Delzon S (2016) Let plant hydraulics catch the waves. J Plant Hydraul. 3:e002.

Fichot R, Barigah TS, Chamaillard S, D Let, Laurans F, Cochard H, Brignolas F (2010) Common trade-offs between xylem resistance to cavitation and other physiological traits do not hold among unrelated *Populus deltoides* x *Populus nigra* hybrids. Plant Cell Environ 33: 1553–1568

Hacke UG, Sperry JS, Pockman WT, Davis SD, McCulloh KA (2001) Trends in wood density and structure are linked to prevention of xylem implosion by negative pressure. Oecologia 126: 457–461

Hartmann H, McDowell NG, Trumbore S (2015) Allocation to carbon storage pools in Norway spruce saplings under drought and low CO2. Tree Physiol 35: 243–252

Harvey HP, Van Den Driessche R (1999) Nitrogen and potassium effects on xylem cavitation and water-use efficiency in poplars. Tree Physiol 19: 943–950

Herbette S, Wortemann R, Awad H, Huc R, Cochard H, Barigah TS (2010) Insights into xylem vulnerability to cavitation in *Fagus sylvatica* L.: phenotypic and environmental sources of variability. Tree Physiol 30: 1448–1455

IPCC (2013) Climate change 2013, the physical basis. Contribution of working group I to the fifth assessment report of the intergovernmental panel on climate change. Cambridge University Press, Cambridge

Jinagool W, Lamacque L, Delmas M, Delzon S, Cochard H, Herbette S (2018) Is there variability for xylem vulnerability to cavitation in walnut tree cultivars and species *(Juglans spp.)*. Hortsci 53: 132–137

Jinagool W, Rattanawong R, Sangsing K, Sévérien Barigah T, Gay F, Cochard H, H., Kasemsap P, Herbette S (2015) Clonal variability for vulnerability to cavitation and other drought-related traits in *Hevea brasiliensis* Müll. Arg. J Plant Hydraul 2: e001

Lamy JB, Bouffier L, Burlett R, Plomion C, Cochard H, Delzon S (2011) Uniform selection as a primary force reducing population genetic differentiation of cavitation resistance across a species range. PLoS One 6: e23476

Lens F, Picon-Cochard C, Delmas CE, Signarbieux C, Buttler A, Cochard H, Jansen S, Chauvin T, Doria LC, Del Arco M, Delzon S (2016) Herbaceous Angiosperms Are Not More Vulnerable to Drought-Induced Embolism Than Angiosperm Trees. Plant Physiol 172: 661–667

Lens F, Tixier A, Cochard H, Sperry JS, Jansen S, Herbette S (2013) Embolism resistance as a key mechanism to understand adaptive plant strategies. Curr Opin Plant Biol 16: 287–292

Martinez-Vilalta J, Cochard H, Mencuccini M, Sterck F, Herrero A, Korhonen JF, Llorens P, Nikinmaa E, Nole A, Poyatos R, Ripullone F, Sass-Klaassen U, Zweifel R (2009) Hydraulic adjustment of Scots pine across Europe. New Phytol 184: 353–364

Martin-StPaul N, Delzon S, Cochard H (2017) Plant resistance to drought depends on timely stomatal closure. Ecol Lett 20: 1437–1447

Pammenter NW, Willigen CV (1998) A mathematical and statistical analysis of the curves illustrating vulnerability of xylem to cavitation. Tree Physiol 18: 589–593

Plavcova L, Hacke UG (2012) Phenotypic and developmental plasticity of xylem in hybrid poplar saplings subjected to experimental drought, nitrogen fertilization, and shading. J Exp Bot 63: 6481–6491

Stojnić S, Suchocka M, Benito-Garzón M, Torres-Ruiz JM, Cochard H, Bolte A, Cocozza C, Cvjetković B, de Luis M, Martinez-Vilalta J, Ræbild A, Tognetti R, Delzon S (2017) Variation in xylem vulnerability to embolism in European beech from geographically marginal populations Tree Physiol. 38: 173–185

Volaire F, Lens F, Cochard H, Xu H, Chacon-Doria L, Bristiel P, Balachowski J, Rowe N, Violle C, Picon-Cochard C (2018) Embolism and mechanical resistances play a key role in dehydration tolerance of a perennial grass *Dactylis glomerata* L. Ann Bot 122: 325–336

Wortemann R, Herbette S, Barigah TS, Fumanal B, Alia R, Ducousso A, Gomory D, Roeckel-Drevet P, Cochard H (2011) Genotypic variability and phenotypic plasticity of cavitation resistance in *Fagus sylvatica* L. across Europe. Tree Physiol 31: 1175–1182.

